# Taxonomy-free fecal microbiome profiles enable robust prediction of immunotherapy response and toxicity in melanoma

**DOI:** 10.1101/2025.11.06.686285

**Authors:** Anastasia Lucas, McKenna Reale, Yuri I. Wolf, Bryant Duong, Yichi Zhang, Jayamanna Wickramasinghe, Lindsey Behlman, Steven M. Jones, Stephanie Higgins, Ahmed M. Moustafa, Abdurrahman Elbasir, Ravi Amaravadi, Tara Mitchell, Alexander Huang, Noam Auslander

## Abstract

The gut microbiome has been causally linked to the efficacy of immune-checkpoint inhibitor therapy (ICI), prompting numerous clinical trials of microbiome-targeting strategies. Yet, mechanisms by which gut microbiota shape immune responses remain elusive as taxonomic biomarkers have failed to generalize across multiple cohorts. In this study, we develop a taxonomy-agnostic framework to identify microbial biomarkers of ICI response and immune-related adverse event (irAE) occurrence from metagenomic sequencing. Applying this approach to four independent melanoma cohorts from clinical centers across the United States, we uncover gut microbial proteins produced by diverse bacterial taxa that consistently predict ICI response. Notably, we uncover a previously uncharacterized operon involved in cellular redox homeostasis that is encoded by different bacteria and reliably predicts irAE occurrence. We further validated the predictive power of this operon in a prospectively sequenced melanoma cohort. Our results demonstrate that taxa-agnostic microbial protein biomarkers are robust, generalizable, and provide a path towards pretreatment risk stratification for melanoma patients initiating ICI therapy.

## Main

Melanoma is the deadliest form of skin cancer with historical survival rates of less than one year for patients with advanced disease^1^. The introduction of immune-checkpoint inhibitors (ICIs) has transformed treatment, raising the 5-year survival rate to over 50%^2^. However, not all patients respond to ICI and rates of toxicity are high, with up to 47%^3–7^ of patients experiencing at least one immune-related adverse event (irAE). irAEs can range from serious, but reversible events like colitis to life-altering immune disorders such as type I diabetes. Severe or life threatening irAEs often require interventions that can limit the clinical benefit of ICI^8,9^ and in some cases require stopping treatment altogether. Further, patients can experience irAEs without benefitting from tumor shrinkage^10^, emphasizing the need for predictive biomarkers that can disentangle response and toxicity to ICI.

One of the most promising emerging strategies for improving ICI outcomes is modulation of the gut microbiome^11–16^. Clinical studies and fecal microbial transplant (FMT) trials have demonstrated that the human gut microbiome can stimulate immune response in non-responding patients^12,17,18^. Efforts to define robust microbial biomarkers to date have largely focused on identifying taxonomic associations with ICI outcomes^19–23^. However, such associations have failed to replicate across studies, institutions, and geographies, likely due to variability in taxonomic composition across populations and other cohort-specific factors^20,24,25^. Mechanisms such as antigenic mimicry leading to autoreactive T cell activation^26–28^ have been suggested to underlie gut microbiome mediated immune responses, but mechanisms by which individual taxa shape ICI-induced immune responses remain poorly understood. This gap in mechanistic understanding has impeded translation of microbiome associations into clinical practice^20^. As such, methods to identify functional microbial biomarkers predictive of ICI outcomes that can translate across cohorts are needed.

Here, we present a taxa-agnostic framework to identify microbial proteins encoded by diverse microbial taxa that are predictive of ICI efficacy and toxicity in pretreatment stool samples from four independent cohorts of melanoma patients across the United States. By combining ICI response-predictive proteins, we constructed a microbial risk score that robustly stratifies patients by likelihood of non-response prior to treatment initiation. We found that several of the proteins best able to predict ICI response are involved in survival mechanisms including nutrient uptake, metabolism, and gene exchange. Notably, we discovered a new irAE-predictive iron-sulfur protein within a previously uncharacterized operon involved in redox homeostasis encoded by multiple different bacteria. We validated the predictive power of this iron-sulfur protein through prospective sequencing of pretreatment stool samples from a cohort of melanoma patients treated with combination PD-1 and CTLA-4 inhibitors, with or without radiation therapy. Beyond discovery of risk-predictive microbial proteins, we present a strategy for prioritization of patient-specific microbial peptides with high MHC-binding potential, paving the way for mechanistic studies and therapeutic targeting. Our findings demonstrate that microbial proteins encoded by multiple taxa are replicable pretreatment biomarkers of ICI response and toxicities, providing a path towards non-invasive clinically actionable risk stratification strategies for melanoma patients.

## Results

### Cohort description

To identify microbial protein biomarkers of ICI benefit in melanoma, we incorporated publicly available datasets along with a prospectively sequenced clinical trial cohort. Public data included four previously published shotgun metagenomic sequencing studies of pretreatment fecal samples from melanoma patients treated with PD-1 inhibitors in the United States; Pittsburgh (n=94)^29^, New York (n=46)^30^, Dallas (n=14)^31^, and Houston (n=25)^21^. Reported response rates in each of the four cohorts were 72% (68/94), 54% (n=25/46), 50% (n=7/14), and 56% (n=14/25), respectively^29^ (Supp. Dataset 1, See Methods). For model development, the Pittsburgh and New York cohorts were used as training sets, Dallas as a validation set, and Houston as an unseen test set.

In parallel to the ICI-response analysis, we analyzed microbial predictors of irAE development. Because irAE annotations were only available in the Pittsburgh cohort, we used this cohort for discovery and tested associations in a prospectively sequenced Philadelphia cohort. The Philadelphia cohort included 36 melanoma patients enrolled in an ongoing RadVax clinical trial (NCT03646617) at Penn Medicine^32^, from whom pretreatment fecal samples were sequenced by shotgun metagenomic sequencing. Within the Pittsburgh cohort, 62% (n=58/94) of patients experienced at least one irAE, while in the Philadelphia cohort 39% (n=14/36) of patients developed an irAE as of the time of study (Fig. 1a, Supp. Dataset 2).

**Fig. 1.**
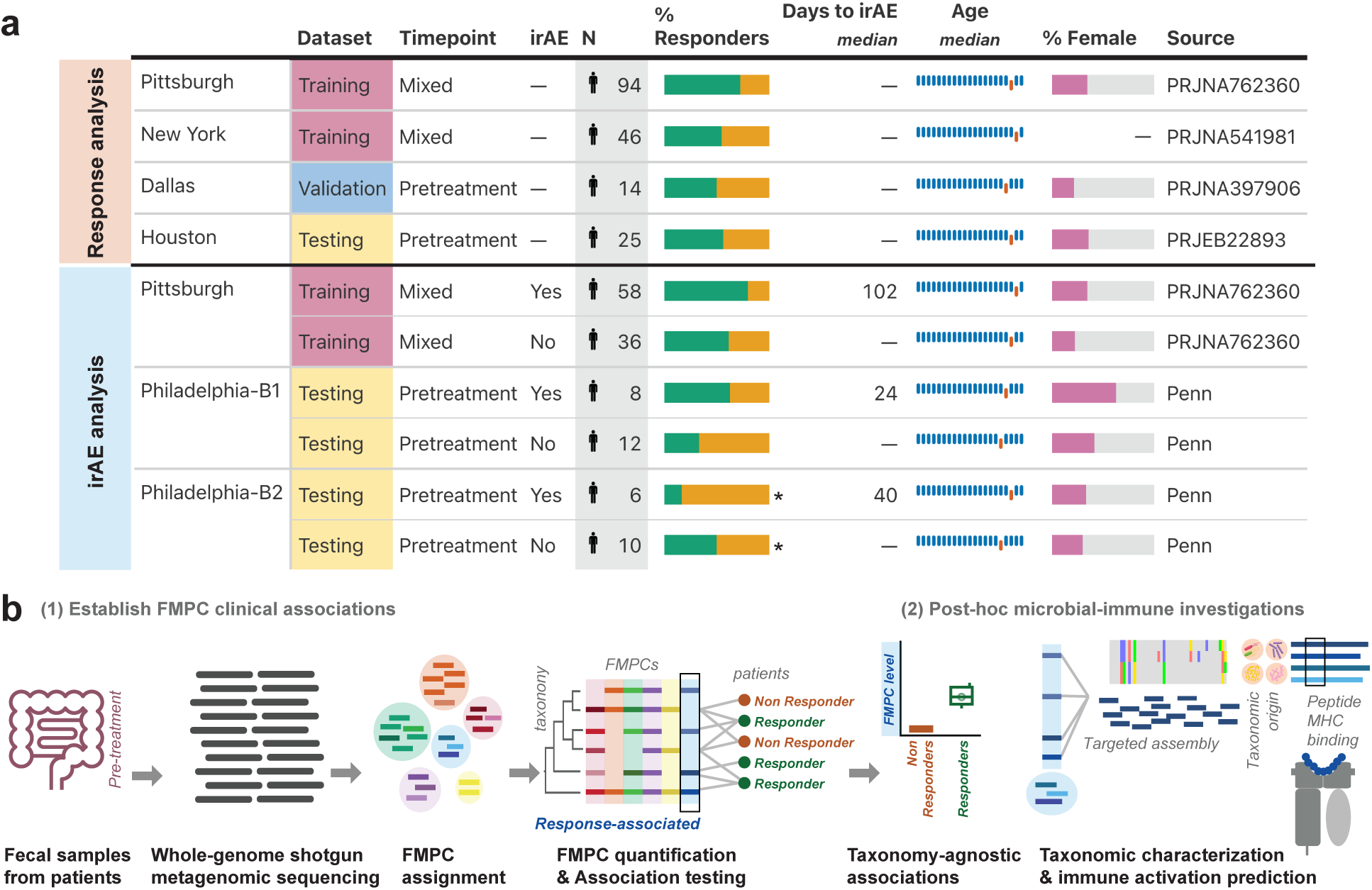
Quantification of FMPCs enables prediction of ICI response across multiple cohorts. **(a)** Demographics and clinical characteristics of cohorts utilized in this study. **(b)** A schematic overview of the computational pipeline. Short read whole genome shotgun sequencing reads are mapped into a clustered database and FMPC-level reads are counted. FMPC abundances are then normalized and correlated with clinical outcomes. In post-hoc analyses, the microbial origin and immune characteristics of ICI-associated FMPCs are evaluated.

### Taxa-agnostic protein clusters enable robust microbial predictors of ICI benefit

Taxa-based biomarkers for ICI benefit have proven difficult to reproduce across cohorts due to bacterial genome plasticity and large geographic variation in gut microbiome composition, even within the same country^20,24,25^. Because biomarker discovery requires replication across clinical centers, taxa-based features are ill-suited for this purpose. In contrast, the functional content of microbiomes is more conserved across humans populations^33^. Moreover, different microbial taxa can encode functionally similar genes, producing homologous proteins that may modulate immune responses in a direct and consistent manner^20,25^. We therefore hypothesized that by quantifying proteins encoded by diverse microbial taxa, we can identify robust, generalizable biomarkers of ICI outcomes.

To this end, we established functionally similar microbial protein clusters (FMPCs) by clustering proteins based on amino acid similarity, following the rationale that sequence homologous proteins share function. We used a recently constructed FMPC reference database generated by clustering non-redundant proteins from RefSeq, yielding 23,951 FMPCs which span diverse bacterial taxa but are functionally homogeneous^34^ (Supp. Dataset 3). We then developed a two-step strategy sensitively map metagenomic reads to FMPCs and subsequently quantify read abundance at the FMPC level (Fig. 1b). First, we aligned DNA reads to RefSeq all non-redundant proteins using BLASTx^35^. Then, we aggregated these reads by FMPCs and normalized to the sequencing depth to generate relative FMPC abundances per sample. Over 95% of read alignments map to a single FMPC, thus resolving the read multimapping, a prevalent source of error in metagenomics^36^, and confirming the validity of this approach (Supp. Fig. 1a).

### Microbial protein clusters consistently predict ICI response across cohorts, improving over taxa-based approaches

We assessed the capacity of the normalized relative abundances of each FMPC to predict ICI response using area under the receiver operating curve (AUROC, Supp. Dataset 4). We used an empirical threshold of AUROC ≥0.65 to select ICI response predictive FMPCs (Supp. Fig. 1b, Methods). Using training and validation datasets for filtering, we identified 49 FMPCs which were significantly associated with ICI response across all (3) cohorts (see Methods, Supp. Dataset 5,6). To compare our FMPC approach to an approach using traditional taxa-based features, we mapped reads to bacterial taxa instead of FMPCs using Bracken^37^, then employed the same AUROC filtering schema as in the FMPC analysis (see Methods for details).

When applied to the held-out test cohort (Houston), the majority of the 49 FMPCs showed consistent predictive trends (Fig. 2a, Supp. Fig. 1c, Supp Dataset 7). Fifteen FMPCs replicated at the empirically selected AUROC ≥0.65 threshold (30.6%) and all 23 of the response-predictive FMPCs showed the same direction of effect in the held-out test cohort (Fig. 2a, Supp. Dataset 8). In contrast, only seven taxa showed consistent predictive capacity among the training and validation cohorts. None of these seven taxa were found to replicate in the held-out test cohort and the direction of effects were less consistent overall (Fig. 2a). These results demonstrate that FMPCs were substantially more consistent across cohorts for predicting response to ICI than taxonomies. To further demonstrate our ability to find consistent FMPC associations across cohorts in different regions of the United States, we performed a principal component analysis on the 49 predictive FMPCs. Our analysis revealed that the responders and non-responders indeed cluster more closely by response status than by cohort of origin (Fig. 2b, non-responders on left and responders on right). We additionally confirmed that several FMPCs enriched in responders compared to non-responders were individually associated with increased progression-free survival (Supp. Fig. 1d,e) Moreover, the number of FMPCs that successfully validated in the held-out test cohort was significant and unlikely to be observed due to chance (p=1e-04, Supp. Fig. 2a-d). Together these results show that our taxa-agnostic approach can achieve significant testing performance and allow for identification of reliable ICI predictors from the gut microbiome.

**Fig. 2.**
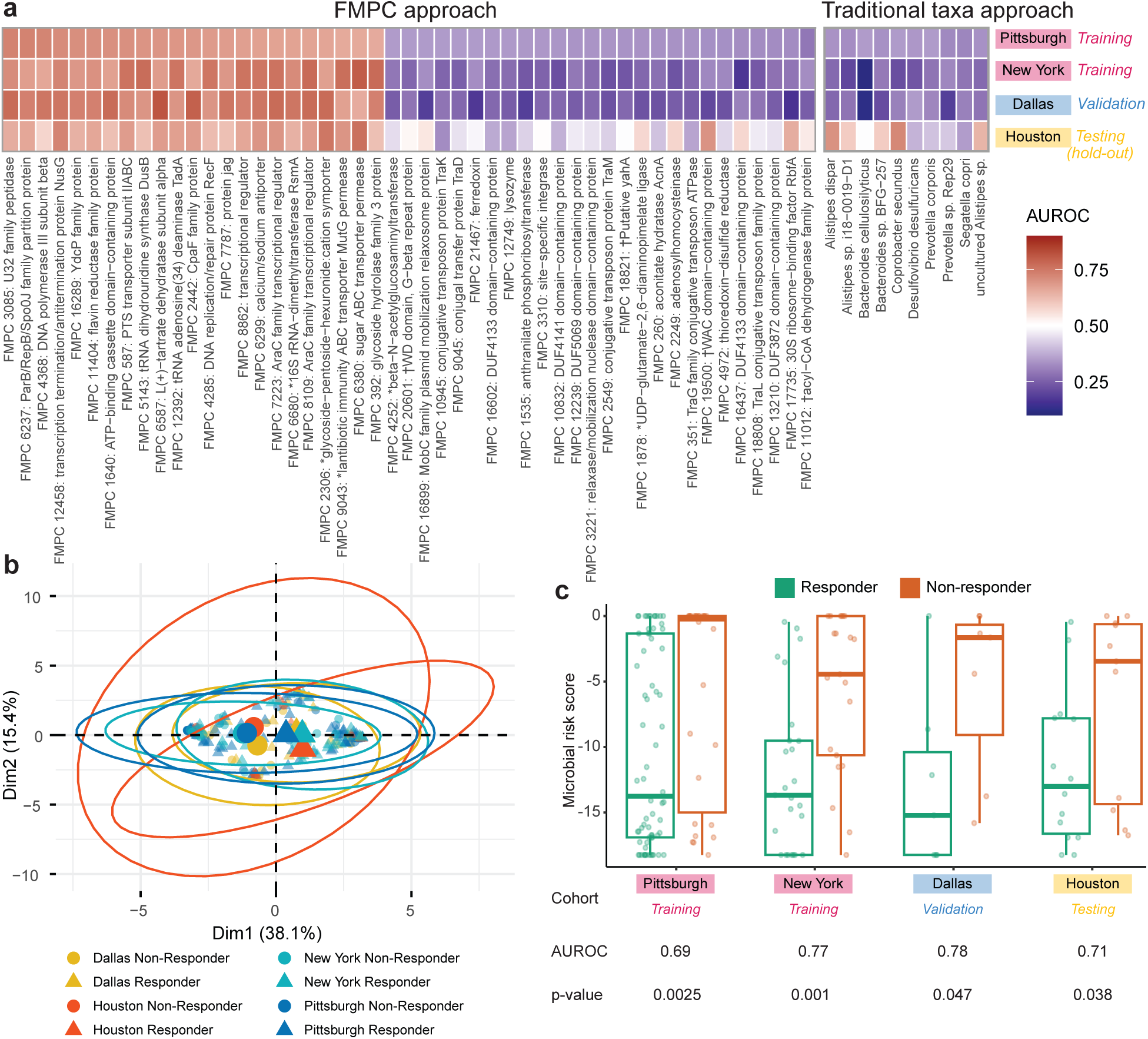
FMPCs allow robust prediction of immunotherapy responses and replication across cohorts, offering an improvement over taxa. **(a)** Heatmap of AUROC evaluating the predictive performance of 49 FMPCs selected based on the training (Pittsburgh and New York) and validation (Dallas) cohorts and tested in an unseen cohort (Houston) (left) compared to AUROCs of taxa identified as predictive in the training cohorts (Pittsburgh, New York, Dallas) with the hold-out test AUROCs (Houston) (right). Asterisks denote truncated FMPC names; daggers denote HHblits41 annotated FMPCs. **(b)** Principal Component Analysis (PCA) of replicating FMPCs in ICI response training and validation cohorts where enlarged points represent group centroids. **(c)** Comparison of microbial risk scores for non-responders (orange) and responders (green) in each cohort.

### FMPC-based patient stratification links metabolic exchange and gene transfer to ICI response

The ability of FMPCs to reliably predict patient response to ICI across cohorts offers a valuable opportunity to assess patient clinical risk through non-invasive methods before ICI treatment initiation. We therefore explored combining the predictive FMPCs to build risk stratification models to score patients based on how likely they are to respond to ICI, thereby identifying patients who are unlikely to benefit from ICI. To this end, we developed an FMPC-level microbial risk score (MRS). Our MRS is motivated by polygenic scores, which measure the cumulative impact of small effects of genetic variants on patient phenotypes^38^. We evaluated several methods for constructing the MRS, including p-value thresholding and restricting to subsets of response- and risk-predictive FMPCs (see Methods, Supp. Fig. 3a,b) to assess which performed the best. Following the trends in the univariate analysis (Fig. 2a), the most generalizable MRS model included all response-predictive FMPCs. The AUROC in the held-out Houston cohort was 0.71, consistent with AUROCs in the training cohorts (Pittsburgh = 0.69, New York = 0.77, Dallas = 0.78, Fig. 2c). Moreover the performance of our MRS surpassed that of traditional polygenic scores used to predict disease risk^38–40^. Several additional MRS models evaluated performed similarly well, suggesting that combining effects of multiple FMPCs could be a robust strategy to assess ICI non-response risk (Supp. Fig. 3a,b).

To increase the explainability of our multivariable ICI response prediction models, we identified the most influential FMPCs through a series of feature selection methods (Methods, Supplementary Methods, Supp. Fig. 3c–e). A RecF DNA replication repair protein and a sugar ABC transporter permease, which were individually associated with progression-free survival in the training cohorts (Supp. Fig. 1d,e), were among the 10 most predictive proteins, suggesting a role for microbial ATP utilization and stress adaptation in modulating therapy outcomes. Consistent with this observation, response-predictive FMPCs were broadly enriched for functions related to membrane transport, nutrient uptake, and metabolism. Multiple FMPCs encoding ABC, PTS, and other transporters, as well as enzymes involved in carbohydrate metabolism and glycan degradation, such as glycoside hydrolases, glycoside-pentoside-hexuronide family uptake proteins, and L(+)-tartrate dehydratase subunit alpha, were enriched in responders (Fig. 2a, Supp. Dataset 8). These findings suggest that metabolic exchanges mediated by the gut microbiome play an important role in shaping ICI response. Further, the strong representation of mobile elements and horizontally transferred genes among these FMPCs may underlie the dissemination of these metabolic capacities across diverse bacterial taxa.

### Pretreatment abundance of a novel operon involved in redox homeostasis predicts immune-related adverse events

After confirming the ability of FMPCs to predict patient ICI responses, we applied our approach to identify FMPCs predictive of irAE development. We identified five FMPCs that were strongly predictive of irAE development in the Pittsburgh training cohort (Fig. 3a,b and Supp. Fig. 4a, Supp. Dataset 9-11). Two of the five FMPCs were short proteins involved in 4Fe-4S binding function, one which was annotated in RefSeq as 4Fe-4S and one which was labeled hypothetical protein (Fig. 3a, Supp. Dataset 9). Both FMPCs had very similar predictive capacity (AUROCs = 0.7 and 0.72), with high relative abundance of 4Fe-4S mapped reads associated with developing an irAE (Fig. 3a,b). Through a remote homology search, we found that the uncharacterized FMPC had a related 4Fe-4S binding domain, explaining the highly consistent abundance between the two FMPCs. The convergence of irAE-predictive FMPCs to 4Fe-4S ferredoxins, together with recent literature reporting ferredoxin microbial derived HLA-bound peptides in melanoma^42^, led us to prioritize the 4Fe-4S FMPCs for further exploration.

**Fig. 3.**
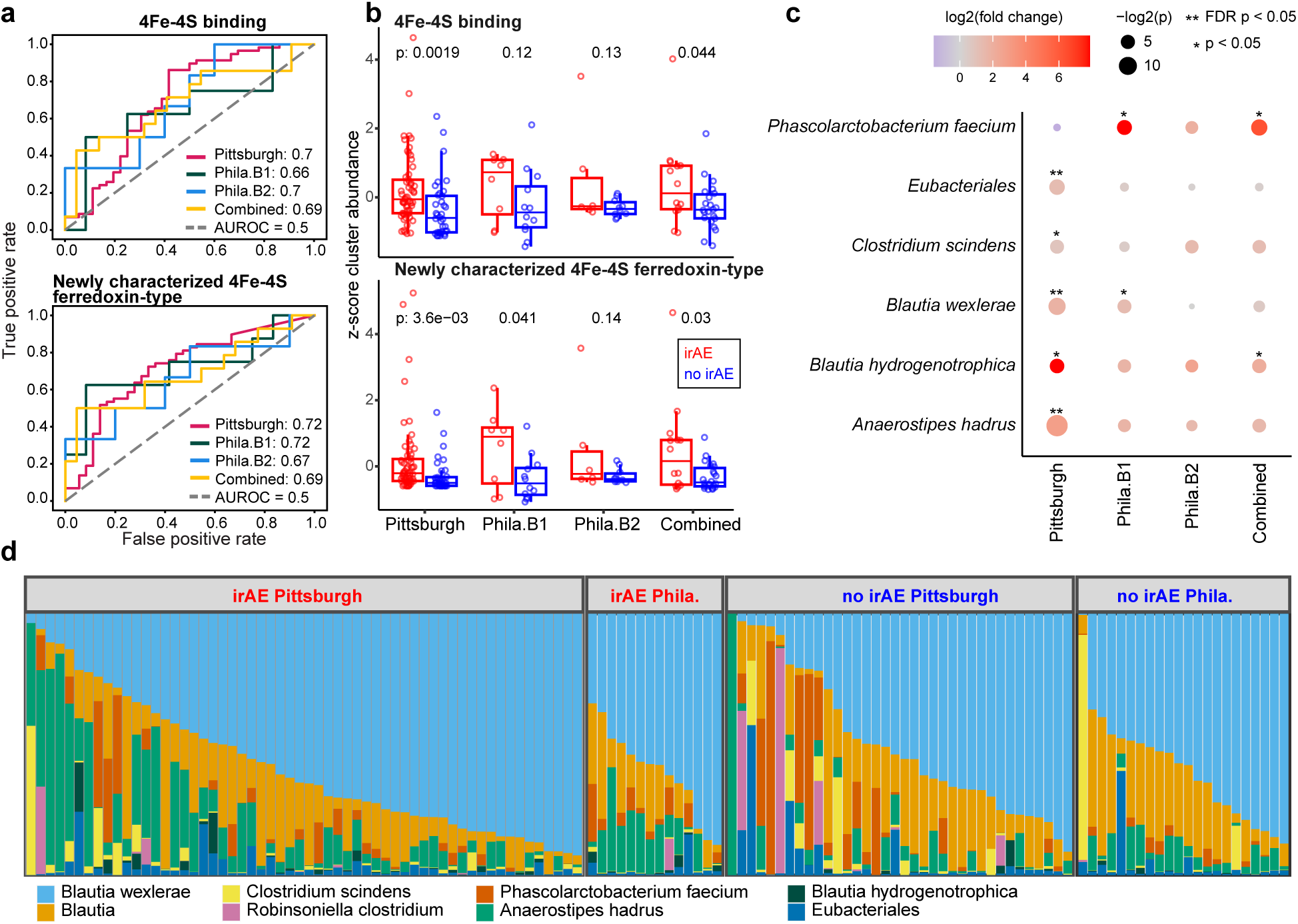
4Fe-4S proteins are predictive of immune-related adverse events. **(a)** AUROC curves for Pittsburgh and Philadelphia cohorts for two 4Fe-4S binding proteins. **(b)** Comparison of normalized relative abundances of the two 4Fe-4S binding proteins between patients with irAEs (red) and patients without irAEs (blue) in each cohort. **(c)** Taxa-specific fold-changes between irAE groups of the 4Fe-4S relative abundances from assembled contigs in each cohort. **(d)** Relative abundances of species mapped to reads aligning to representative contigs containing the 4Fe-4S protein in each cohort.

We validated the 4Fe-4S associations in a prospective setting through additional data collection. We established a Philadelphia cohort of melanoma patients treated with ICI at Penn Medicine with pretreatment (day 0) stool samples. We first tested the 4Fe-4S FMPC association with irAE in a pilot batch of Philadelphia patients (n = 20). We observed the two 4Fe-4S FMPCs yielded AUROCs of 0.66 and 0.72, respectively (Fig. 3a,b). We next performed sequencing on the remaining patient samples in a second batch of Philadelphia patients (n = 16). Again, we observed a similarly high predictive capacity for the two 4Fe-4S FMPCs with AUROCs of 0.7 and 0.67 in the second group of patients. The associations between relative abundance of 4Fe-4S proteins and irAE development also held when the two batches were combined (AUROCs = 0.69 and 0.69, respectively; Fig. 3a,b).

We next performed a comprehensive characterization of the irAE predictive 4Fe-4S FMPCs. To investigate the bacterial taxa that encode these proteins, we performed a targeted assembly of reads mapping to the 4Fe-4S FMPCs to build contigs containing the 4Fe-4S regions (see Methods for detail, Supp. Dataset 12). Alignment of the contigs revealed that the irAE-predictive 4Fe-4S proteins are encoded by multiple bacterial species, including six from the class *Clostridia* and one from the phylum *Firmicutes* (Supp. Dataset 12). Furthermore, the assembled contigs of these regions from different bacteria indicated the two 4Fe-4S FMPCs were in fact two domains within one protein of a three-protein operon. Each operon was found to contain a 4Fe-4S ferredoxin, a Thioredoxin-like (TlpA) family disulfide reductase protein, and a CxxC binding protein (Supp. Fig. 4b-g). All three proteins contain cysteine-rich motifs that coordinate metal ions or form disulfide bonds, enabling electron transfer reactions crucial for maintaining cellular redox homeostasis. It is possible that the proteins encoded by this operon shift microbial redox metabolism, influencing the production of immunomodulatory metabolites and reactive oxygen species. These changes may disrupt mucosal immune tolerance and promote pro-inflammatory signaling, contributing to autoimmune activation in genetically susceptible hosts.

A major motivation behind our FMPC quantification approach is to identify ICI phenotype-associated microbial features common to different species that may be obscured when performing functional analyses predicated on taxa. When comparing taxa-specific irAE associations of the region encoding the 4Fe-4S FMPCs, we found the effect sizes and strengths of association varied between the Pittsburgh and Philadelphia cohorts (Fig. 3c). Relatedly, there were distinct differences in the taxa-specific relative abundances of the 4Fe-4S protein region between the Pittsburgh and Philadelphia cohorts. While both cohorts are dominated by *Blautia wexlerae*, less abundant species exhibited greater variability. For example, 4Fe-4S proteins originating from *Phasolarctobacterium faecium* were found to be more prevalent in the Philadelphia cohort while *Anaerostipes hadrus* 4Fe-4S was more abundant in the Pittsburgh cohort (Fig. 3d). We also observed significant differences in strength of taxa-specific 4Fe-4S associations between the Pittsburgh and Philadelphia cohorts (Fig. 3c,d), supporting the notion that functional level associations predicated on taxa are insufficient to find reliable signal across cohorts. Therefore, our FMPC approach meets an important need by enabling identification of microbial features that reliably predictive of ICI outcomes across different clinical centers.

### 4Fe-4S peptides have clinical potential for determining pretreatment immune-related adverse event occurrence

To demonstrate how the 4Fe-4S peptides could be developed into a non-invasive clinical test for risk of irAE development, we extracted DNA sequences that could be used as a clinical qPCR for 4Fe-4S protein detection. We first selected bacteria where the encoded 4Fe-4S proteins were closest to the FMPC sequences used for irAE prediction. We then used the translated proteins from assembled DNA sequences to identify conserved regions across the bacterial strains which align to each of two 4Fe-4S FMPCs (See methods for details, Fig. 4a and Supp. Fig. 4b-d). In both cases, the protein sequences encoded by DNA associated with *Anaerostipes hadrus* were most similar to the identified 4Fe-4S irAE-predictive FMPCs. We therefore extracted the DNA encoding these conserved regions from *Anaerostipes hadrus* to form candidate PCR sequences. To evaluate the utility of these sequences for irAE prediction in clinical settings, we quantified reads aligning to these sequences in both the Pittsburgh and Philadelphia cohorts. Both sequences were found to be significantly associated with irAE development in the two cohorts, supporting a potential biomarker for testing in the clinic (Fig. 4b).

**Fig. 4.**
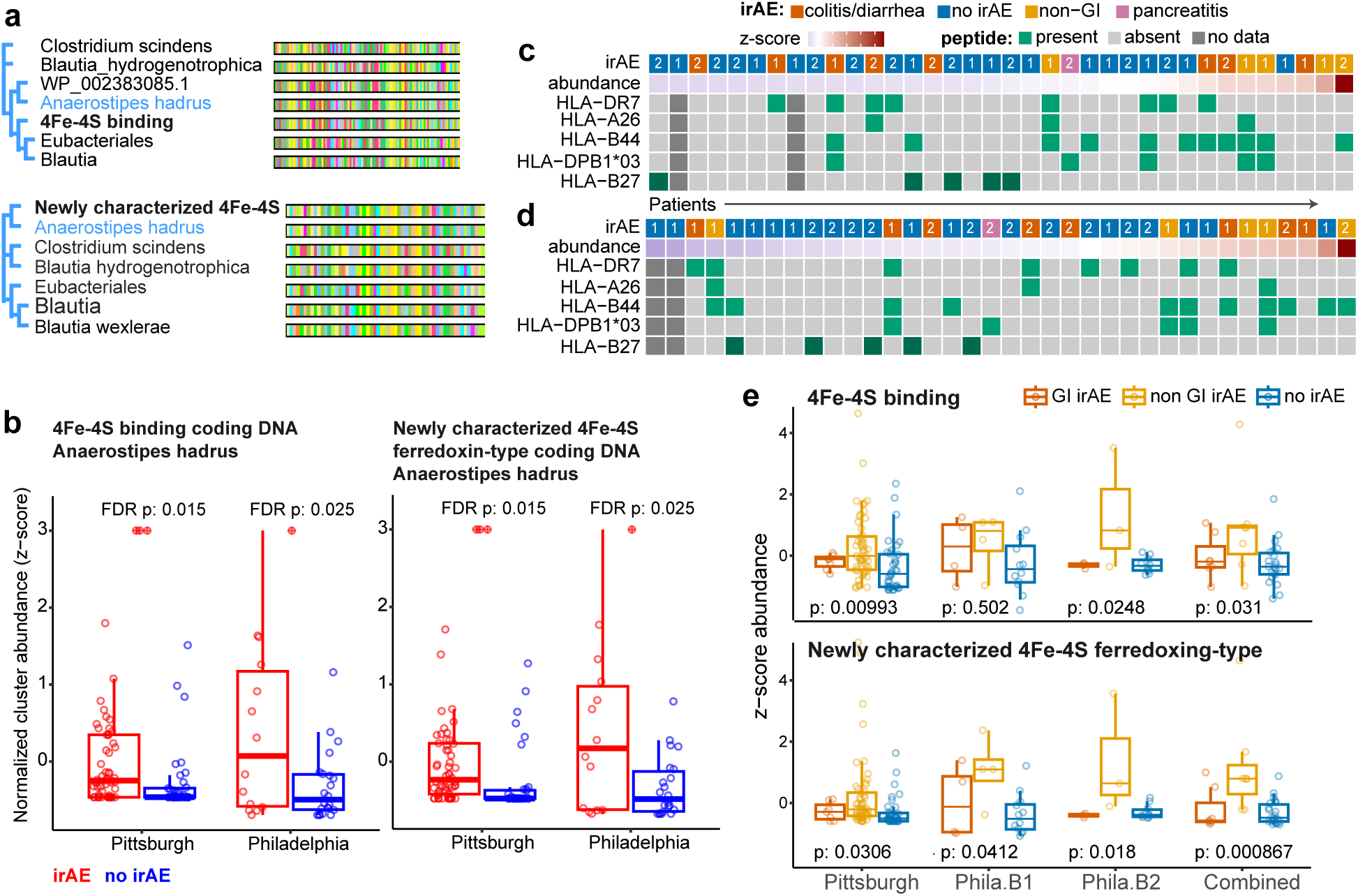
Potential immune mechanisms linking the 4Fe-4S FMPCs to melanoma irAEs. **(a)** Protein alignment and tree of the conserved sequence of the two 4Fe-4S FMPCs through coding frames of assembled contigs. **(b)** Comparison of normalized relative abundances of two candidate PCR nucleotide sequences, derived for the two 4Fe-4S irAE-predictive FMPCs between patients with irAEs (red) and patients without irAEs (blue) in each cohort. The x-axis is truncated for visual clarity and outlier values with z-score ≥3 are denoted with a (+) symbol. **(c)** HLA and irAE patterns of in patients with high and low relative abundance of the newly characterized protein containing the 4Fe-4S binding domain. **(d)** HLA and irAE patterns in patients with high and low relative abundance of the 4Fe-4S protein. **(e)** Differences in relative abundance of the 4Fe-4S proteins between patients without irAE, patients with non-GI irAEs (including pancreatitis), and patients who developed colitis or diarrhea.

### Genetic susceptibility and microbial ferredoxin levels explain organ-system specific immune-related adverse events

The gut microbiome is closely intertwined with the immune system and gut microbes can elicit immune responses via different mechanisms in the context of ICI^21,23,43^. Because MHC molecules can bind to and present bacterial peptides recognized by T cells^44^, we explored the possibility of 4Fe-4S peptide recognition by MHC class I molecules. We performed HLA serotyping for all patients in the Philadelphia cohort, which was paired with the metagenomic sequencing data. Evaluating potential MHC antigen presentation of microbial peptides associated with irAE (Methods and Supplementary information) showed that overall binding affinities did not differ by irAE status, although multiple strong binders were present in patients with irAEs.

To identify potentially immunogenic peptides, we developed a framework (Supp. Fig. 5a) to extract patient-specific peptides from regions containing the Fe-4S, TlpA, and CxxC proteins and predict their binding potential using NetMHCpan-4^45^. Many individual peptides had strong predicted binding affinities in individual patients; however, we did not observe clear patterns in the binding predictions of any specific peptide with irAE status across cohorts (Supp. Fig. 5b,c). Our findings suggest that peptide immunogenicity is unique rather than ubiquitous and experimental studies should prioritize peptides for testing in a patient specific manner.

Because some HLA types have been suggested to increase susceptibility to irAEs^46–48^ we additionally sought to determine whether there were HLA associations with irAE development both independent of and in the context of 4Fe-4S levels. Overall, the strongest HLA type association with irAE development independent of 4Fe-4S levels was with DR7 (unadjusted p = 0.049). In the first Philadelphia batch, we observed that the DRB1*07/DQA1*02 haplotype was enriched in patients with GI-related irAEs (75%) compared to those without irAEs (10%) and compared to a general population (Supp. Fig. 5d). Although this haplotype has not been previously implicated in ICI-induced colitis and diarrhea, there is some evidence to suggest a general association with gastrointestinal (GI) conditions^49^. We also observed that all patients with irAE in the first Philadelphia batch with below-median levels of one of the 4Fe-4S FMPCs had the DRB1*07 type; conversely, all irAE patients without DRB1*07 had above-median 4Fe-4S levels. Given the strength of the DRB1*07 associations, we proposed these might be complementary mechanisms. The trend between DRB1*07 status and 4Fe-4S levels remained consistent, albeit not perfectly correlated, after integrating the second Philadelphia batch (Fig. 4c,d). Still, motivated by the DRB1*07 association, we examined GI-specific trends in the 4Fe-4S FMPCs associations. Interestingly, we found that 4Fe-4S FMPC relative abundance levels were substantially more associated with non-GI irAE events in both the Pittsburgh and Philadelphia cohorts (unadjusted p = 9.9e-3, 0.031, 0.031, 8.7e-4, respectively, Fig. 4e). In contrast, 4Fe-4S FMPC relative abundance levels were similar in patients with GI related irAEs, i.e. diarrhea and colitis, and those without any irAE events. Looking more broadly, we noticed distinct patterns in 4Fe-4S abundance levels with additional HLA types (Fig. 4a). For example, we observed patients with HLA-B44, which has been previously associated with PD-1 efficacy^50^, commonly had irAEs and tended to have higher abundance of 4Fe-4S proteins. Conversely, none of the Philadelphia patients with HLA-B27, which is known to modulate the microbiome,^51,52^ developed irAEs and tended to have lower than median levels of the 4Fe-4S protein (Fig. 4c,d). Given numerous reported associations between HLA and irAEs^46,47,53^, additional research is needed understand how interplay between HLA and the microbiome could lead to development of irAEs within specific organ systems.

## Discussion

Recent preclinical and clinical studies have demonstrated that the gut microbiome can modulate tumor immunity and responses to ICI^21,54^. Although mechanisms underlying the influence of gut microbiota on ICI response are poorly understood, bacterial metabolites, anti-microbial peptides and immune cell priming have been proposed to intrude antitumor immunity^54,55^. Consequently, microbiome-elicited immune responses have been mostly attributed to microbial products or functions. Based on the knowledge that different taxonomies share similar microbial products and functions, in this study we established a framework to systematically investigate taxa-agnostic microbial features predictive of ICI response and irAE development.

Our taxa-agnostic approach to gut microbial biomarker discovery enables identification of microbial functions associated with ICI risk across multiple cohorts throughout the United States. We identified FMPCs involved in horizontal gene transfer, nutrient uptake and carbohydrate metabolism, cell wall synthesis, redox and central metabolism which consistently predict ICI response in independent cohorts of melanoma patients enrolled in studies at different clinical centers. Many of these protein functions are highly related to bacterial survival mechanisms and stress response, falling in line with the well-documented correlation between gut microbiome diversity and PD-1 response in melanoma patients^55,56^. Combining these response-predictive FMPCs enabled models that stratified patients by risk of ICI non-response, addressing a key challenge in predicting which patients are most likely to benefit from ICI therapy. Early identification of patients who are at high risk for non-response or irAE can guide monitoring across the course of treatment, allowing for early intervention to improve patient outcomes. Despite using different methods of combining FMPCs (univariate, MRS, random forest) our protein-based models were robust across cohorts, suggesting that protein-based models can provide broadly applicable risk prediction strategies.

We also identified a novel three-protein operon containing a 4Fe-4S protein that is associated with non-GI related irAEs. We confirmed the association between the 4Fe-4S-containing operon and irAE occurrence in a newly sequenced Philadelphia melanoma cohort. This operon was found to be encoded across multiple microbial taxa, with association strength varying by taxonomic origin, highlighting the value of taxa-agnostic microbial features for biomarker identification. Many of the peptides derived from this operon were predicted to be immunogenic in individual patients, suggesting a potential role of molecular mimicry (Supp. Methods). However, experimental validation would be needed to confirm the immunogenic potential of any patient-specific microbial peptides. In addition to the likely function of this 3-protein operon in enabling electron transfer reactions crucial for maintaining redox homeostasis, 4Fe-4S proteins also have 3-4 transmembrane segments. This finding raises the possibility that these thioredoxin disulfide reductase proteins are reducing extracellular or cell-surface targets to enhance immune recognition. Given the known role of thioredoxin-like proteins in MHC class I peptide loading and pathogen recognition^57–59^, this operon may cooperate to promote antigen processing in the gut.

We acknowledge that our study has limitations. Our training and validation layout demonstrated the importance of including multiple cohorts for discovery of predictive microbial features and we anticipate the robustness of FMPC findings would increase as more training cohorts are included. Well-powered cohorts remain a challenge in the field, especially for irAEs studies as irAEs are highly heterogeneous in nature (Fig. 4c). Differences in study design between cohorts may also be a limiting factor for our study, although these differences reflect real world scenarios for metagenomic biomarker discovery and application. Additionally, because we used a lenient approach for clustering proteins, we expect some larger FMPCs may be noisy. We therefore prioritized controlling for false positive associations over mitigating false negatives by performing post-hoc validations. As such, we expect additional ICI response- and irAE-predictive FMPCs could be found by additional fine-tuning of the underlying cluster-building approach. Furthermore, we report correlative relationships between FMPCs and ICI outcomes and experimental validation is required to glean insight into potential mechanisms of action for specific protein or peptide sequences.

In summary, our study demonstrates the power of direct functional profiling of the microbiome to enable pretreatment clinical risk stratification and identify non-invasive biomarkers for melanoma patients receiving immunotherapy. Beyond melanoma, our taxa-agnostic provide a versatile framework for uncovering functional microbial risk factors across diverse cancers and other human health outcomes.

## Methods

### Patient characteristics

Shotgun metagenomic sequencing and clinical data from three independent cohorts of melanoma patients who received PD-1 inhibitors were downloaded from the Sequence Read Archive. The three cohorts were comprised of patients from three clinical centers in Pittsburgh^29^, New York^30^, and Dallas^31^, where fecal samples were processed independently according to their respective study protocols. Overall clinical response was available for all three cohorts and irAE information was annotated for the Pittsburgh cohort^29^. Although some cohorts included multiple sample timepoints, we included the complete cohorts to allow sufficient sample size of this training cohort and mitigate spurious AUROC assignments due to the small sample set used in filtering (Supp. Dataset 1).

For prospective testing of identified predictors of irAE, we additionally established a Philadelphia dataset with melanoma patients enrolled in a RadVax clinical trial at Penn Medicine^32^ (n = 37), in which we performed whole-genome shotgun metagenomic sequencing and human leukocyte antigen (HLA) serotyping. All available pre-treatment fecal samples from Philadelphia patients (n = 36) were sequenced via Illumina NovaSeq and demultiplexed. Nextera-XT adapters were removed using trimmomatic-0.33. Nucleotide quality for each position was averaged across all reads using FASTQC and low-quality reads as defined by Trimmomatic were discarded. Metagenomic sequencing was performed in two batches, with the first containing samples from 20 patients and the second containing samples from 16 patients (Supp. Dataset 13). HLA typing was performed using human peripheral blood monocytes from 37 Philadelphia patients with available tissue, including 34 patients with paired metagenomic sequencing (Supp. Dataset 13). As most patients experienced at least one irAE, we restricted the irAE group to patients who experienced one or more Grade III, IV, and V irAEs.

### Defining and quantifying FMPCs

We quantified FMPCs using a two-step approach. (1) FMPCs establishment. We used blastdbcmd to download non-redundant bacterial proteins (taxid 2) on August 11, 2022. We clustered 43,174 WP proteins in this database using CD-HIT^60^ with the similarity threshold set to 40% and a word length of 2. This process resulted in 23,951 FMPCs, which were named by the CD-HIT cluster representative annotation or through homology search using HHblits^41^. (2) FMPCs quantification. We aligned shotgun metagenomic sequencing data to microbial proteins in the NCBI representative nr database (Supp. Dataset 3) using BLASTX and extracted alignments with pident > 80, alignment length (sequence overlap) > 20, and e-value < 1e-8. Using the filtered alignment results, we aggregated reads at the FMPC level and normalized the read counts the total number of reads sequenced per million per sample.

### Identifying FMPCs which predict ICI response

We assessed the ability of the normalized abundance counts to separate phenotype classes, i.e. responders vs. non-responders or patients with irAEs vs. patients without irAEs, using area under the receiver operating characteristic curve (AUROC). We considered FMPCs as being predictive of a phenotype class if their AUROCs were ≥ 0.65, chosen empirically based on the distribution of all AUROCs in the largest training cohort (Pittsburgh, Supp. Fig. 1b), as it roughly translates to 0.01 percentiles of AUROCs on both tails. Overall, we identified 565 predictive FMPCs in the Pittsburgh training cohort, of which 134 were also predictive in the New York training cohort. A subset of 86 were verified as predictive in the Dallas validation cohort (Supp. Dataset 5). To further restrict the FMPCs carried for testing, we set an additional criterion of having ≥ 0.7 average AUROC across the three cohorts, yielding 49 FMPCs for test evaluation (Supp. Dataset 5). Note that for brevity we refer to AUROCs < 0.5, i.e. those which are predictive of the phenotype class coded as 0 by their complement.

To assess statistical significance of each FMPC’s AUROC values, we employed a permutation testing procedure for the 49 selected proteins. We fit each protein’s relative abundance to a probability distribution using fitdistrplus^61^. We simulated 5,000 random datasets with the same number of cases and controls as the training data for each protein using the estimated distributions. We conducted a one-sided permutation test to find the probability of obtaining AUROCs at least as extreme as the observed AUROC by chance. A one-sided test was selected as the FMPCs were already required to replicate in the same direction in multiple cohorts. FMPCs with a false discovery rate-adjusted permutation p-value < 0.05 were considered significant. To assess the significance of the number of replicating FMPCs in the hold-out test cohort, we permuted the response labels of the Houston dataset 10,000 times and recomputed the AUROC for each of the 49 significant FMPCs. We computed a one-sided p-value to find the probability of obtaining 15 or more replicating FMPCs.

To identify sets of FMPCs that could predict ICI response, we constructed a random forest model using a predefined set of 10 FMPCs selected from an initial set of FMPCs using a genetic algorithm (Supp. Methods). Because we used the Dallas cohort to evaluate the fitness of our genetic algorithm, our initial set of FMPCs was based only on the Pittsburgh and New York cohorts using the criteria outlined above (≥ 0.65 in individual training cohorts and average AUROC ≥ 0.7 in both training cohorts). Additional details on the model generation can be found in Supplemental Methods.

### Comparison between FMPC and taxa-based approaches

To determine whether FMPCs provide advantages over traditional taxa-based approaches, we repeated our AUROC evaluation process. We used abundances assigned to distinct bacterial taxonomies through read counts reported by Bracken^37^. To assign reads to taxa, we ran Kraken2 and Bracken with default parameter settings and the Standard-8 database. We normalized the raw Bracken counts by sequencing depth and computed reads per million for every detected taxa. As in the FMPC analysis, we identified significant taxa by computing AUROC and performing the selection criteria detailed above using the different training cohorts to extract predictive taxonomies. Statistical significance testing was performed using the permutation methods described in the FMPC section, yielding a final set of taxa features that was evaluated on the unseen Houston test cohort.

### Microbial risk score construction

Microbial risk scores (MRS) were computed using a method analogous to polygenic scores, by summation of *p* FMPC’s individual effect sizes, β_i_, for each FMPC present at ≥ median abundance in the patient’s sample:

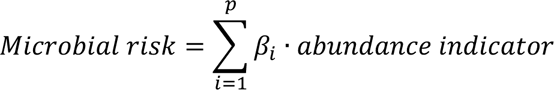

The abundance indicators refer to binarized abundances (1 = higher than median abundance), as described above, and were used for the risk scoring in lieu of the real valued abundances to improve generalizability and clinical interpretation. β_i_ was computed for each of the predictive FMPCs as determined by AUROC were computed using the Pittsburgh training cohort by running logistic regression models for which the dependent variable was binary coded treatment response, and the independent variable was normalized abundance for a single protein; proton-pump inhibitor (PPI) use at the time of sample collection was included as a covariate. PPI use, along with other clinical variables (antibiotic use, sex, body mass index, and sample timepoint) were prior assessed for their effect on treatment response using multiple logistic regression. PPI use was the only variable found to be significant (p = 0.042) and as such was singularly selected as a covariate. A regression framework was chosen for MRS as it allows for the inclusion of both positive- and negative-directed effects as opposed to a burden framework, which has previously been used for taxa-based risk scoring^62^. We used combinations of two separate FMPC filtering approaches to compute the MRS: 1) using either all, risk-, or response-predictive proteins in the training set after adjusting for PPI use and 2) retaining FMPCs with non-significant p-values (p ≥ 0.05) when adjusted for PPI use or no thresholding. We observed that the model including only the 23 response predictive proteins without p-value thresholding was the most robust across cohorts, although several other models performed similarly well (Supp. Fig. 3a,b). Differences in microbial risk scores between responders and non-responders for each cohort were assessed using AUROC and Wilcoxon rank-sum tests. The Pittsburgh cohorts were used for training the MRS weights, the New York and Dallas cohort were used for model validation, and Houston cohorts was used as test data.

Additional analyses showed that the AUROC for the hold-out test set varied depending on which approach was used to construct the model (Supp. Fig. 3a,b) and that the p-value thresholding step often decreased model performance. As statistical significance is a function of sample size, thresholding may be especially important when the sizes of the training cohorts are smaller, which is common in metagenomic studies in comparison to genetic studies, which often include tens of thousands of patients.

### Identifying the bacterial species encoding 4Fe-4S FMPCs

To identify longer contigs of DNA regions encoding the 4Fe-4S FMPCs that predict irAE, we first extracted all nucleotide reads that aligned to the 4Fe-4S FMPC in the Pittsburgh cohort using BLASTX with the parameters described previously. We used SPAdes^63^ to assemble the 4Fe-4S mapped reads into longer contigs. The contigs assembled by SPAdes were then extended in both forward and backward directions using an iterative greedy assembly algorithm. In each iteration, starting with the assembled SPAdes contig, the first (prefix) and last (suffix) 55 nucleotides of the contig were extracted as a query sequence. A string search was performed across all fasta files corresponding to samples from the Pittsburgh cohort for each suffix and prefix to identify possible extensions in either the forward or backward directions, respectively. The contig was then extended by the longest sequence that appeared in ≥ 4 patients’ sequencing data and new suffixes and prefixes were extracted from the extended contig. This process was repeated for 20 iterations or until there was no extension that appeared in at least 4 patients. Using these representative contigs, we determined their taxonomic identities using Blast against core_nt and refseq, where the most highly scored taxonomies assigned with e-value < 1e-6 consistently using both datasets were selected for each contig.

Bowtie2^64^ was applied to align reads against the contigs encoding the identified regions from each species identified to quantify the specific taxonomies associated with this region across patients in the Pittsburgh and Philadelphia cohorts. All alignments with maximum of 1 mismatch, seed substrings of length 25, and maximum fragment length of 2,000 were reported by Bowtie2.

We used MetaGeneMark^65^ to perform gene prediction on the representative contigs for each species. We verified the presence of the 4Fe-4S gene as well as determined that it always occurred alongside a TlpA disulfide reductase gene with, in all instances but one, a Cxxc binding domain gene in between; in one instance we observed 4Fe-4S and TlpA with the Cxxc not in between the two (Supp. Dataset 12). The taxonomic identities of the representative contigs were identified using blastn and verified with blastp of the respective MetaGeneMark proteins. If the contigs and proteins aligned to multiple species at similar coverage and percent identity, we labelled the contig at the higher taxonomic level shared by these sequences.

### Predicted binding for patient-specific peptides

With the goal of identifying peptides that may bind to specific MHC presented in patients, we used the Bowtie2 alignment output to the assembled contigs representing the newly identified region from different taxonomies. To construct patient-specific peptides, SPAdes assembly was run on the Bowtie2 output of every patient using *k*-mer sizes of 21, 33, 55, and 77 without read error correction and with the --careful flag enabled to reduce the number of mismatches and short indels. To find common peptides among patients, 6-frame translation was performed to all patient-specific contigs assembled by SPAdes, removing any amino acid sequences that contained a stop codon. All possible 15-mers of the amino acid sequences were extracted and 30 15-mers that appeared in ≥ 60% of all patients were selected for further interrogation.

NetPanMHC-4.1^45^ was used to predict binding affinities of the 30 selected peptides. We specifically looked at the binding affinities for HLA types that were overrepresented in patients with or without irAEs as well as peptides most likely to bind in individuals regardless of frequency of occurrence. (Supp. Dataset 14). We additionally verified that each of the selected peptides unambiguously aligned to bacterial 4Fe-4S or TlpA genes using blastp.

### Establishing 4Fe-4S DNA sequences for a potential PCR test

The two 4Fe-4S FMPCs found to predict irAE represent short amino acid sequences which were not annotated in known bacterial species. To establish DNA sequences reflecting these FMPCs for a potential PCR test, we first evaluated which proteins in the assembled contigs contained sequences homologous to the two 4Fe-4S FMPCs. In all cases, the two 4Fe-4S FMPCs were homologues to two distinct domains of the 4Fe-4S proteins in assembled contigs. Therefore, to derive DNA sequences for quantification, we first searched for the contig encoding protein sequences with greatest homology to the identified FMPC. We focused on the most conserved regions across all taxa identified to encode these FMPCs (Supp. Fig. 4c,d). Using a phylogenetic tree constructed for multiple sequence alignments of the conserved regions aligning to the two FMPCs, we found that the sequence annotated as *Anaerostipes hadrus* encoded the regions with highest homology to the two FMPC (with 96% and 99% amino acid identity, Fig. 4a, Supp. Fig. 4c,d). Therefore, the respective DNA sequences in encoding these regions in the contig annotated as *Anaerostipes hadrus* were extracted and used as a Bowtie2 index (172 and 294 nt, Supp. Dataset 15), to quantify reads aligning against each DNA sequence as a potential PCR test (Fig. 4a).

### Statistical analyses

For follow-up analysis of highly associated protein clusters, patients were grouped into high/low abundances for a protein based on whether their abundance counts were greater than or equal to the median abundance across all samples in the cohort for that protein. Survival analysis was performed in R using the survival::survfit() function to model progression free survival (days or months, depending on cohort) and log-rank p-values testing the difference in progression-free survival for high and low abundance groups were computed using survival::survdiff(). Data was censored based on if the patient had a progression event. The Dallas cohort was excluded from survival analysis due to data unavailability. Differences between two group means were computed using t-tests. Differences among more than two groups were tested using one-way ANOVA and post-hoc pairwise differences were conducted using Tukey HSD. Fold-change analyses were computed using a one-sided Mann-Whitney U test comparing the median percent of reads mapping to the representative contig for the patients who developed irAEs and patients who did not develop irAEs for each species. *Blautia hydrogenotrophica* was excluded from fold-change analysis due to sparsity of data. HLA analyses were performed using Fisher’s exact tests. Normal population frequencies for HLA alleles reflect the 1000 Genomes European superpopulation^67^ accessed through 1000 Genomes FTP.

## Data availability

Microbiome sequencing data with de-identified metadata will be deposited to NCBI SRA upon acceptance of this manuscript. Processed microbiome data (normalized counts), de-identified HLA typing, and clinical data are available through the supplementary information. Access to publicly available shotgun metagenomic sequencing data was obtained through SRA, with accessions PRJNA762360 (Pittsburgh), PRJNA541981 (New York), PRJNA397906 (Dallas) and PRJEB22893 (Houston).

## Code availability

Code developed for this study is available through GitHub: https://github.com/AuslanderLab/melanoma-ICI-FMPC-analysis

## Patient and the Public Involvement Statement

Patients or the public were not involved in any way.

## Acknowledgements

This research was funded by NIH grants R00 CA252025, P50 CA261608, R01 LM014503, F31 LM014962, V foundation award V2024-006, and Michelson Medical Research Foundation. Y.I.W. is supported by the Intramural Research Program of the National Institutes of Health (National Library of Medicine). The authors would like to thank Michael Galperin (CBB/NLM/NIH) for providing insights on possible mechanisms and interpretations; and Kavitha Sarma (Wistar), Maureen Murphy (Wistar), Andrew Patterson (University of Pennsylvania), Julia Malnak for feedback on the manuscript.

