## Supplementary Information for "Taxonomy-free fecal microbiome profiles enable robust prediction of immunotherapy response and toxicity in melanoma"

**Supplementary Datasets**

**Supplementary dataset 1** – Clinical information for publicly available cohorts utilized in this study, Pittsburgh and New York (training), Dallas (validation) and Houston (test).

**Supplementary dataset 2** – Clinical information for the RadVax cohort (this study)

**Supplementary dataset 3** – FMPC database providing protein IDs, cluster information and functional description for each FMPC

**Supplementary dataset 4** – Normalized FMPC counts for the training and validation cohorts

**Supplementary dataset 5** – AUROCs assigned to each FMPC for predicting immunotherapy response in the training and validation cohorts

**Supplementary dataset 6** – Permutation testing results for the 49 selected FMPCs for prediction of immunotherapy response based on the training and validation cohorts.

**Supplementary dataset 7** - Houston test dataset (for immunotherapy response prediction) normalized FMPC abundances for the 49 FMPCs tested for response prediction

**Supplementary dataset 8** – Immunotherapy response prediction AUROCs across the cohorts for the 49 selected response-predictive FMPCs

**Supplementary dataset 9** – AUROCs predicting irAE for each FMPC in the Pittsburgh cohort

**Supplementary dataset 10** – RadVax prospective testing cohort (for irAE prediction) normalized abundances of 5 tested irAE-predictive FMPCs

**Supplementary dataset 11** – AUROCs for irAE prediction for the 5 selected irAE FMPCs across Pittsburgh and RadVax cohorts, with adjustment for covariates.

**Supplementary dataset 12** – Assembled regions of bacteria encoding the identified 4Fe-4S irAE predictive FMPCs

**Supplementary dataset 13** – HLA typing of patients in the RadVax cohort

**Supplementary dataset 14** – Peptide MHC binding prediction results for the RadVax cohort with paired HLA typing

**Supplementary dataset 15** – 4Fe-4S DNA sequences derived for quantification as a proxy for a qPCR test

**Supplementary Figures**

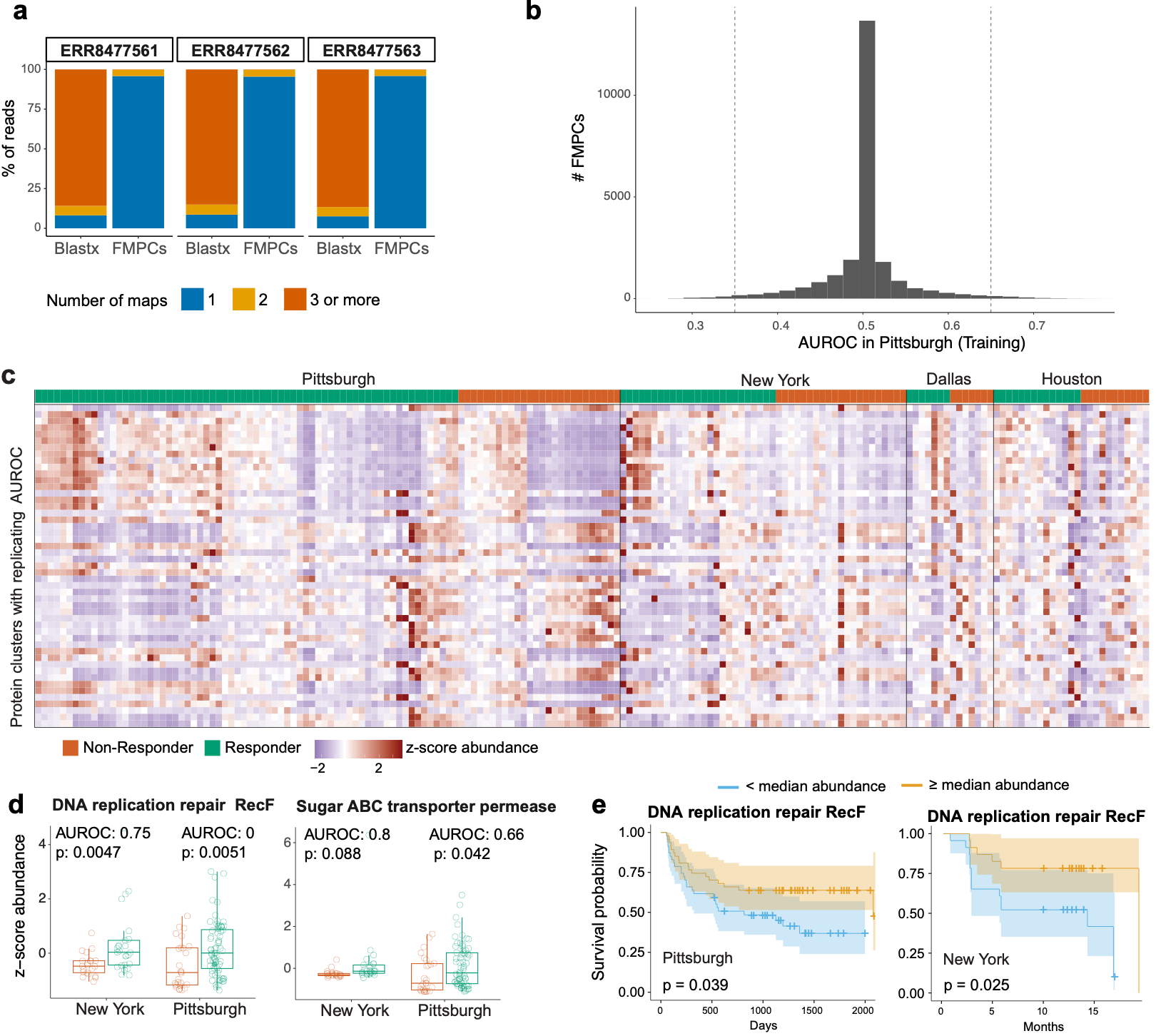

**Supplementary Fig. 1.**

**(a)** The reduction in ambiguous maps when looking at read-to-FMPC maps compared to read-to-protein level hits (BLASTX) for 3 representative samples.

**(b)** Histogram of FMPC AUROC distribution in the Pittsburgh training cohort

**(c)** Heatmap showing z-score transformed normalized abundances of protein clusters with AUROCs ≥ 0.65 annotated with clinical response, non-responders (NR), orange and responders (R), green.

**(d)** Boxplots showing normalized FMPC abundances for responders and non-responders for two representative ICI response associated FMPCs.

**(e)** Kaplan-Meier curves showing progression-free survival for patients ≥ or < median normalized FMPC abundances.

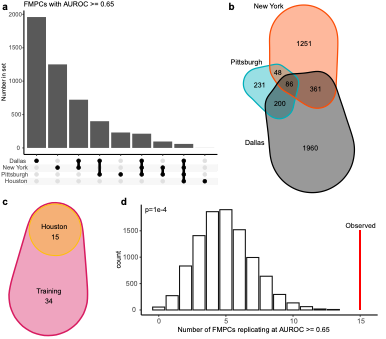

**Supplementary Fig. 2.**

**(a)** UpSet plot showing intersecting sets of response-predictive FMPCs across the training, validation and hold-out test cohorts. For the Houston test cohort, only 49 predictive FMPCs selected based on the training and validation data were evaluated.

**(b)** Venn diagram showing the number of intersecting response-predictive FMPCs across the training cohorts at an AUROC threshold of ≥ 0.65. Note that FMPCs that had AUROC ≥ 0.65 for predicting response and AUROC ≥ 0.65 for predicting risk in a different cohort would be considered as significant in individual cohorts, but not in the intersection. To be included in cohort intersections, the FMPC must be significant in the same direction.

**(c)**  Venn diagram showing the number of FMPCs replicating in the hold-out test cohort (Houston) of replicating FMPCs out of the final set of 49 FMPCs.

**(d)** Null distribution for the number of replicating FMPCs out of 49 obtained by randomly permuting the response labels for the Houston hold-out test cohort 10,000 times and recomputing the AUROC for each FMPC. The observed number of replicating FMPCs is shown in red.

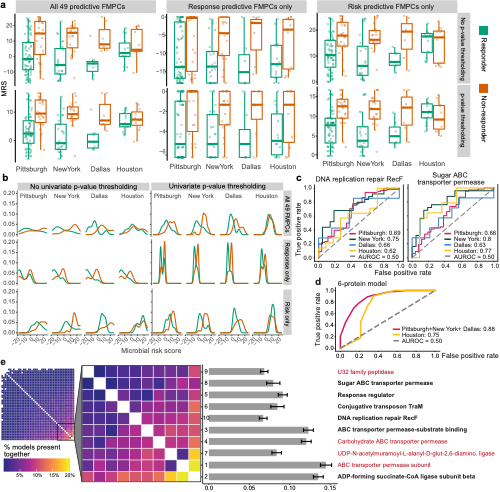

**Supplementary Fig. 3.**

**(a)** Boxplot comparisons of patient microbial risk scores constructed using different FPMCs, i.e. all FMPCs, response predictive, or non-response (Risk) predictive sets using p-value thresholding (PRT) or no thresholding (Full).

**(b)** AUROC curves for predicting ICI responses based on microbial risk scores constructed using different FMPC sets as described in (a).

**(c)** AUROC curves for individual predictive FMPCs and **(d)** the 6-FMPC random forest model in all cohorts.

**(e)** A subset of FMPCs tend to be present together in best performing models selected from 10,000 runs of a genetic algorithm. The FMPCs that occur in the modes together most frequently are annotated by their feature importance scores in the 10-protein ICI response prediction random forest model averaged across 1,000 seeds. Black text denotes the 6 FMPCs included in the exploratory model that was tuned to the Houston cohort.

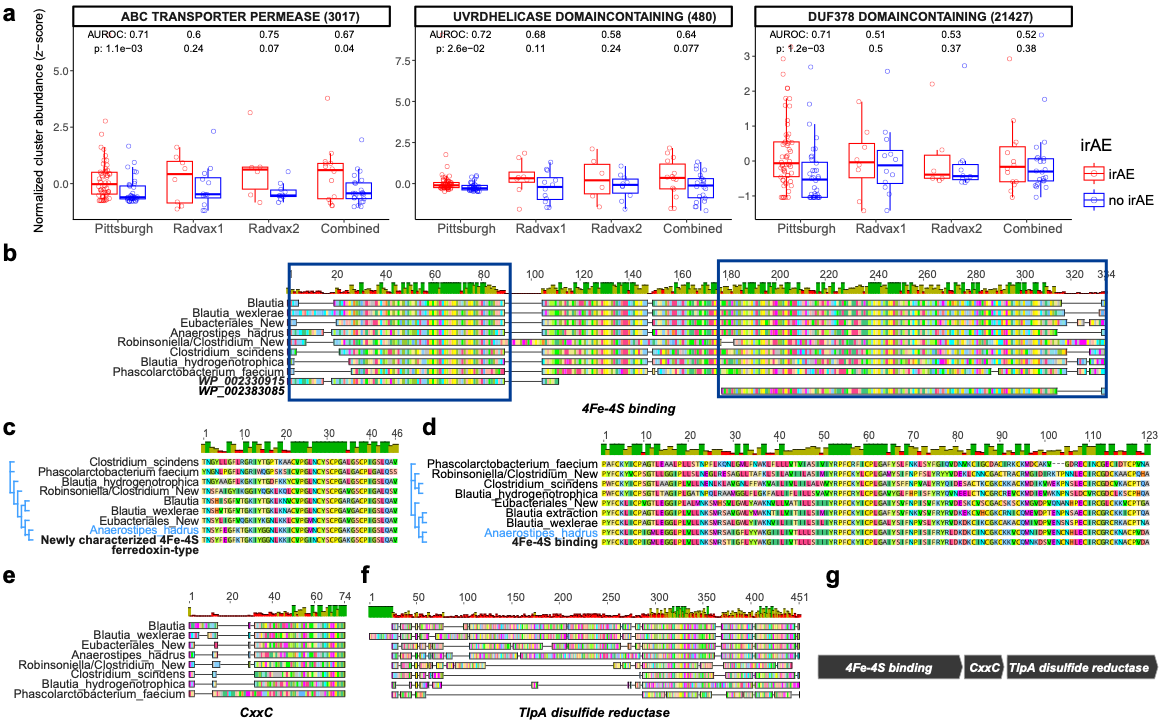

**Supplementary Fig. 4.**

**(a)** Comparison of normalized relative abundances of three non-4Fe-4S irAE predictive proteins with AUROC > 0.7 in the Pittsburgh cohort between patients with irAEs (red) and patients without irAEs (blue) in each cohort.

**(b)** Protein sequence alignment of frames encoded by the assembled regions, which map to the two 4Fe-4S FMPC, when aligned with the two FMPC representatives. The two 4Fe-4S FMPCs align to different region of the 4Fe-4S proteins found in different bacteria through assembly.

**(c)** Protein sequence alignment and a phylogenetic tree based on a conserved region mapping to the leftmost FMPC from (b), which is the newly characterized 4Fe-4S FMPC. *Anaerostipes hadrus* 4Fe-4S sequence is closest to the FMPC sequence for this region.

**(d)** Protein sequence alignment and a phylogenetic tree based on a conserved region mapping to the rightmost FMPC from (b), which is the characterized 4Fe-4S FMPC. *Anaerostipes hadrus* 4Fe-4S sequence is closest to the FMPC sequence for this region as well.

**(e)** Protein sequence alignment of frames encoded by the assembled regions which map to the CXXC motif-containing proteins, likely present on the same operon as the 4Fe-4S proteins to which the irAE associated FMPC map.

**(f)** Protein sequence alignment of frames encoded by the assembled regions mapping to the TlpA family protein disulfide reductases, which is likely present on the same operon as the 4Fe-4S proteins to which the irAE associated FMPC map.

**(g)** A schematic showing the likely new operon identified, in which two regions within the 4Fe-4S proteins were strongly associated with irAE. The operon has shorter CXXC motif-containing proteins confined within 4Fe-4S and TlpA family protein disulfide reductases for all region assembled except one. The exception contig, likely originating from *Phascolarctobacterium faecium*, encodes the three proteins in a different order, whereby 4Fe-4S is confined between the CXXC motif-containing protein and the TlpA family protein disulfide reductase.

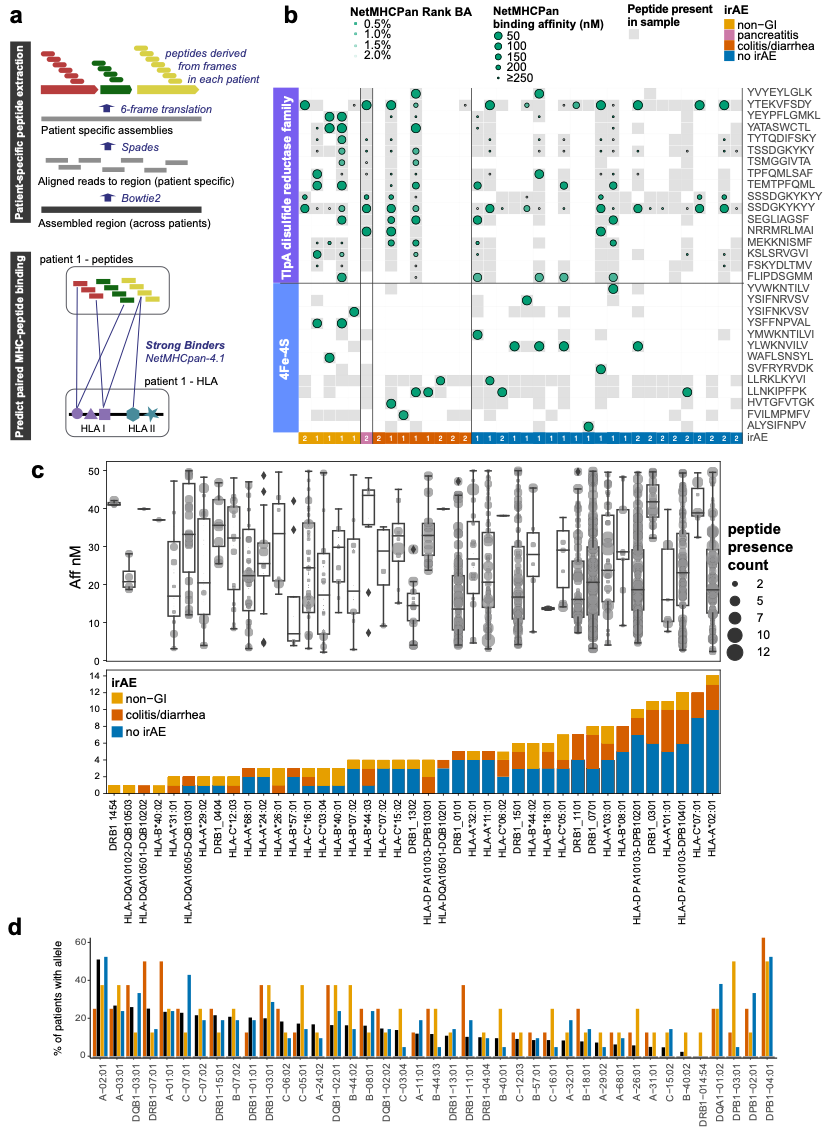

**Supplementary Fig. 5.**

**(a)** A schematic overview of the pipeline developed to identify patient specific MHC binding of microbial peptides associated with irAE.

**(b)** MHC Binding affinities predicted by NetMHCpan-4, of peptides derived from the 4Fe-4s and TlpA coding frames in Philadelphia patients by different irAE subgroups. Patient specific HLA typing was paired with peptides derived for each patient for prediction.

**(c)** Lower panel – HLA typing prevalence (sample count) in the RadVax cohort across 3 clinical categories (non-GI irAE, GI-irAE and no irAE). Upper panel – boxplots and overlaid dot-plots showing the binding affinities of patient-specific irAE associated peptides, which strongly bind to each HLA type through NetMHCpan-4 (Aff_nM<50). Dot size corresponds to the number of patients in which each strongly binding peptide is found.

**(d)** Percent of patients with HLA allele in RadVax compared to 1000 Genomes European population.

**Notable ICI response FMPC associations**

Whereas none of the FMPCs perfectly discriminated between responders and non-responders, similar enrichment patterns were observed in subsets of patients across the cohorts (Supp Fig. 1c). For example, a subset of responders showed an overabundance of Sugar ABC transporter permease (Supp Fig. 1d). Additionally, higher than median relative abundances of reads mapping to these FMPCs was also associated with higher progression-free survival in both the Pittsburgh and New York cohorts (Supp Fig. 1e). In general, we found that different FMPCs captured different sets of responding patients, and these were generally consistent across cohorts (Supp Fig. 1c,f and Supp Fig. 2a,b). We therefore investigated whether we could improve predictive capacity by combining multiple FMPCs to predict response to ICI.

**Additional multivariate FMPC models**

Upon observing the success of combining FMPCs into a single risk score, we sought to identify an FMPC model that could improve prediction of ICI response. We therefore trained a random forest model using a preselected set of 10 FMPCs using a genetic algorithm (see below). The performance of the model on the held-out Houston set was moderate (AUROC = 0.61, not shown) and similar to the performance observed for some individual FMPCs (Supp Fig. 3c). However, due to the strength of the overall trend in the univariate analyses and the success of our MRS, we investigated whether any subset of the 10 FMPCs could improve predictive performance in the Houston cohort. Through exploratory stepwise analysis, we identified a 6 FMPC model that optimized performance on the Houston cohort (Supp. Fig. 3d, AUROC = 0.88 for the training and validation cohort, and 0.75 for the Houston cohort). Notably, both the full and reduced models contained RecF DNA replication repair and sugar ABC transporter permease proteins, which demonstrated individually strong association with ICI response and associations with progression-free survival in the training cohorts (Supp. Fig. 1d,e). Overall, the results from the random forest model and the MRS support the utility of combining different FMPCs for ICI response prediction and patient risk stratification. The co-occurrence and mutual exclusion of these FMPCs in high scoring models (Supp. Fig 3e) suggest it could be worthwhile to investigate combinatorial or complementary effects of FMPCs to refine these multi-FMPC models in future work.

**Additional FMPC associated with irAEs**

Given that the 4Fe-4S association was validated in a prospective, independent cohort, we investigated whether the other three irAE predictive FMPCs identified in the training cohort could be validated in the Philadelphia cohort. Of the three remaining FMPCs, an ABC transporter permease FMPC showed a similar, albeit weaker and less consistent trend with AUROC = 0.6, 0.75, and 0.67 in the first batch, second batch, and combined batches, respectively (Supp Fig. 3a). Another FMPC representing UvrD helicase binding proteins replicated in the first Philadelphia batch (AUROC = 0.68), but not did not meet the pre-defined threshold in the second (AUROC = 0.58). The fifth FMPC, labeled DUF378 domain-containing, was not found to be predictive of irAE in either Philadelphia batches (Supp Fig. 3a, Supp Datasets 10,11). Overall, only a few FMPCs were predictive of irAE development, and these tended to be more consistent than those identified as predictive of ICI response. The 4Fe-4S which was tested prospectively had the most consistent association overall.

**Peptide prioritization using NetMHCpan-4**

We performed a prioritization of peptides with immunogenic potential by extracting patient-specific microbial peptides and predicting their MHC affinity using NetMHCpan-4^44^ using paired patient-derived peptides and HLA genotypes (Supp. Fig. 5b). This allowed an evaluation of possible peptide-MHC affinities across different categories of patients in the Philadelphia cohort. We found many patient-specific peptides that were predicted to be strong MHC binders based on the patients’ HLA types (Supp. Fig. 5c). We did not observe substantial differences in binding affinities in patients based on irAE status; however, multiple peptides were predicted as strong binders HLA types that were common among patients in the Philadelphia cohort. (Supp. Fig. 5d). Along with the increased abundance of genes encoding these proteins in patients with irAE (Fig. 3b,c), these predictions suggest an elevated occurrence of peptides bound to MHC through bacteria encoding 4Fe-4S protein clusters.

**Supplementary methods**

**Training, validation and testing design**

Cohorts with WMS sequencing of Melanoma patients treated with ICI within the United States were considered through this study. For FMPC prediction of ICI response, cohorts used in the Pittsburgh study were utilized. Training, validation and testing cohorts were designated in advance, where the largest and most comprehensive cohort (Pittsburgh) was used for training for both response and irAE prediction. In our initial setup, the Chicago cohort was designated as part of the training set. However, we found a complete discordance between the Chicago and the rest of the training cohorts. Since we could not find patient characteristics and clinical study design for the Chicago cohort (in contrast to other cohorts considered), the Chicago cohort was excluded from consideration in this study.

The Pittsburgh cohort was the only publicly available cohort with patient irAE information. We performed WMS for the RadVax cohort after designating specific FMPCs to be tested based on the Pittsburgh training cohort, and therefore deem this a prospective validation of the irAE FMPCs. Briefly, DNA was extracted from approximately 200 mg of stool using the Qiagen DNeasy PowerSoil Pro kit. Extracted DNA was quantified using the Quant-iT PicoGreen dsDNA assay kit (Thermo Fisher Scientific) before library generation. Shotgun libraries were generated from 7.5 ng DNA using Illumina DNA Prep Library Prep kit and IDT for Illumina unique dual indexes at 1:4 scale reaction volume. Library success was assessed by Quant-iT PicoGreen dsDNA assay and samples with library yields < 1 ng/ul were re-prepped as needed. After all samples for a given pool were prepped, an equal volume of library was pooled from every sample and then the pool was sequenced using a 300 cycle Nano kit on the Illumina MiSeq. Libraries were then repooled based on the demultiplexing statistics of the MiSeq Nano run. The quality of final libraries were checked on the Agilent BioAnalyzer for the size distribution and absence of additional adaptor fragments. Libraries were sequenced on an Illumina Novaseq 6000 v1.5 flow cell, producing 2x150 bp paired-end reads. Extraction blanks and nucleic acid-free water were processed along with experimental samples to empirically assess environmental and reagent contamination.  A laboratory-generated mock community consisting of DNA from *Vibrio campbellii* and Lambda phage were included as a positive sequencing control.

**Feature selection for the random forest model using genetic algorithms**

Using a genetic algorithm, we generated 10,000 random forest models where the validation cohort (Dallas) was used to evaluate fitness using an initial population size of 20 and feature size 5, over 9 generations. The AUROCs of the validation data from these 10,000 models ranged from 0.64-0.93. We then used a subset of 10 protein clusters that frequently co-occurred in the 10,000 models to generate a random forest model for predicting ICI outcome. Although the final model performed well on the validation set (training AUROC=0.94, validation AUROC = 0.99), it did not generalize as well to the hold-out testing set (AUROC = 0.61) (Fig. 1b). Given that the performance was better than random and given our observation that a greater number of cohorts improve the generalizability of the FMPCs (Supp Fig. 2a,b), we explored whether a post-training feature removal strategy could improve the performance overall. To this end, we iteratively removed protein clusters from the random forest model until the AUROC could not improve using the hold-out testing set as a validation set and all other cohorts as a training set. To allow using an integrated set with multiple datasets to train the random forest models, we first coded each FMPC as a binary feature indicating whether the sample had above or below median relative abundance in comparison to other samples in the cohort, instead of using the normalized relative abundance counts. Yet, none of the trained models allowed a good generatability on an unseen test set.

**HLA comparison**

HLA data from 1000 Genomes was obtained from <http://ftp.1000genomes.ebi.ac.uk/vol1/ftp/data_collections/HLA_types/>. Allele frequencies reported as sample counts from EUR populations (n = 529) were used for comparison to a general population as all RadVax patients were self-reported white/non-Hispanic. Ambiguous calls from 1000 Genomes data were not included.

**Molecular mimicry investigation**

Our analysis identified NP_002487.1/NDUFS8 as a human encoded homolog of the 4Fe-4S bacterial protein associated with irAE. However, we identified only ACPVDAI is a possible shared human-bacterial peptide, by aligning against human proteins using BLASTP.
